# Extensive loss of forage diversity in social bees due to flower constancy and communication in simulated environments

**DOI:** 10.1101/2023.11.01.565092

**Authors:** Christoph Grüter, Francisca Segers, Lucy Hayes

## Abstract

Bees require a diverse diet for a healthy development. Many bee species show flower constancy, that is, they visit flowers of just one species during a foraging trip. Flower constancy is important for plant reproduction, but it could impair dietary diversity in bees, especially in biodiversity-depleted, human-modified landscapes. It is assumed that flower constancy does not lower dietary diversity in social bees, such as honey bees or bumble bees, because different colony members can specialise on different plant species. However, this has never been tested. We used computer simulations to investigate the effects of flower constancy on colony diet in plant species-rich and species-poor landscapes. We also explored if communication about food sources, which is used in many social bees, further reduces forage diversity. Our simulations reveal an extensive loss of forage diversity due to flower constancy in both plant species-rich and species-poor environments. Small colonies often discovered only 30-50% of all available plant species, thereby increasing the risk of nutritional deficiencies. Our simulation results could explain why bumble bees, which have small colony sizes, are less flower constant than honey bees and stingless bees, which have larger colony sizes. Remarkably, when colonies also communicated about food sources, *Simpson’s diversity*, which measures the evenness of flower visits, approached near zero in plant species-poor environments. Finally, we found that food source clustering, but not habitat fragmentation impaired dietary diversity. These findings can help in the design of landscapes that increase forage diversity and improve bee nutrition and health in human-modified landscapes.

## Introduction

Bees are essential pollinators of wild and agricultural plants (1–3), owing to their abundance, and their morphological and behavioural diversity (4, 5). Another key reason for their value as pollinators is their diet. Many bees have a broad (*i.e.* polylectic) diet at species level, but as individuals they specialise on one flower species during a foraging trip, so-called flower constancy (6–10). Flower constancy is beneficial for plant fitness as it reduces conspecific pollen loss and the negative impacts of heterospecific pollen deposition (7, 10–13). However, the causes and consequences of flower constancy from the bees’ perspective remain less well understood (7, 9, 14). One benefit of flower constancy may be that it avoids the time and cognitive load associated with having to learn how to exploit multiple flower species efficiently (7–10, 14–16). However, it remains a puzzling behaviour as both individual bees and different bee species vary in the degree of flower constancy (9, 10, 17, 18).

Bees require a balanced diet to maintain basic biological functions (19–23). Pollen, in particular, plays a key role in bee health as the main source of protein, lipids and micronutrients (19, 20, 22). An inadequate supply of protein (and essential amino acids) has been shown to impair body size (24–26), lifespan (27, 28) and ovary development (27) in both social and solitary bees. Pollen from different plant species vary greatly in their macro- and micro nutrient content (21, 22, 29, 30), and a diet based on a small number of pollen types risks a surplus of some nutrients and a deficiency in others (31), with negative impacts on reproductive success (32) and survival (28, 33, 34) in *Bombus terrestris* and *Apis mellifera*. Accordingly, there is increasing evidence from different bee species that a more diverse pollen diet has health benefits (35), for example, by boosting larval size (36), immunocompetence (37) and the survival of bees infected by viruses or parasites (30, 33, 38). Flower constancy, which can last several days (39, 40), could negatively impact dietary diversity and exacerbate the effects of biodiversity loss in strongly human-modified environments, e.g. agricultural landscapes. Foraging challenges in modern landscapes are suspected to be a key driver of poor bee health as environments lacking in floral diversity make it harder for bees to achieve their intake targets for important nutrients (22, 23, 30, 31, 41–43). Evidence for the combined effects of environmental change and narrow dietary preferences comes from observations showing the bumblebees with a narrow diet were more likely to decline in numbers in the last decades (44). Yet, if and how flower constancy affects dietary diversity under different ecological circumstances remains unknown.

Social bees, mainly the honey bees (Apini), bumble bees (Bombini) and stingless bees (Meliponini), are often highly flower constant (17, 45). It is assumed that flower constancy does not negatively affect forage diversity in social bees because different individuals can specialise in visiting different flower species, thereby ensuring that the colony exploits a range of plant species (14, 46, 47). While social bee colonies indeed collect pollen from many plant species, often only ∼1-5 pollen types are collected in larger amounts at any given time (19, 48–51), possibly risking nutritional deficiencies. Colony size could be a key factor mediating the effects of flower constancy on colony forage diversity as a larger foraging workforce could potentially discover a wider diversity of plant species than a smaller one. Another social trait with potential implications for forage diversity is recruitment communication. Many social bees communicate about profitable food sources (52–54), *e.g.* the honey bee waggle dance, excitatory runs in combination with buzzing sounds in bumble bees and stingless bees, or trophallaxis in honey bees and stingless bees (52, 53, 55, 56). The function of these diverse behaviours is to direct nestmates towards profitable food sources, often by transmitting olfactory information that allows recruits to identify the advertised plant species in the surrounding environment (52, 57, 58). Recruitment communication may reduce colony forage diversity because it causes colonies to focus on a subset of the available food sources (55, 59, 60).

Our understanding of the consequences of flower constancy and communication on colony diet breadth remains limited, firstly, because it is not usually possible to manipulate flower constancy while keeping other factors constant and, secondly, because it is logistically challenging to perform flower constancy experiments at a landscape scale. Agent-based simulation models are powerful tools to circumvent these obstacles and provide insights into how social and ecological factors interact with foraging strategies to modify emergent colony-level properties (14, 61–65). We developed an agent-based simulation model to study if and how flower constancy and communication affect the diversity of plant species collected by a colony in both plant species-rich and species-poor environments. Within these environments, we manipulated food source distribution (uniform *vs*. clustered), abundance and reward size. In addition, we created fragmented landscapes, *e.g.* representing an urban habitat, to explore how habitat fragmentation affects forage diversity with a view to help in the design of conservation strategies which could improve bee nutritional diversity in human-modified landscapes. We measured both *Species diversity*, *i.e.* the number of plant species a colony discovers, and *Simpson’s diversity*, which also takes into account the evenness of plant species exploitation (66). We predicted that colony size, plant species richness, and food source distribution (clustered *vs.* uniform) determine the effects of flower constancy and communication on dietary diversity. Finally, we performed a literature search to assess the degree of flower constancy in bees with different social lifestyles to aid in the interpretation of our simulation results.

## Materials and Methods

### Agent-based model

We built an agent-based model (ABM) using the programming software NetLogo 6.1 (67) (see NetLogo files with full model code: https://doi.org/10.5281/zenodo.8320942). The model builds on an earlier version (14, 61), from which it differs in key aspects, such as the types of data that were collected, the way food sources were distributed, their characteristics, and the number of flower species in the environment. The model simulates a bee colony surrounded by food sources (see below for descriptions of food sources and Table S1 for default and alternative values tested). The model does not simulate a particular bee species but is built to resemble a bumble bee or a *Melipona* stingless bee colony in terms of colony size, flight behaviour, and communication mode. The bees operate on a two-dimensional square grid with 400 x 400 patches. A single patch length corresponds to 5 meters. Thus, the size of the virtual world corresponds to 2 x 2 km, which covers the typical foraging distances of most bee species (61, 68, 69) (Fig. S1). Each simulation lasted 36,000 seconds (*i.e.* 10 hours in total), representing a day with good foraging conditions.

### Simulated forager bees

Colony sizes ranged from 10 to 300 bees, which covers the typical forager workforces of many bumble bee and stingless bee species (70, 71). Bees began the simulation in the centre of the nest as *generalists* (Fig. S2). They then moved at a flying speed of 1 patch/second (*v*_flight_), a flight speed similar to that of bumble bees (5m/sec) (72), following a Lévy-flight pattern (73, 74). A Lévy-flight is a random sequence of flight segments whose lengths, *l*, come from a probability distribution function having a power-law tail, *P*(*l*)∼*l*^-μ^, with 1<μ<3 (with *μ* = 1.8 as our default) (74). After agents encountered a food source, they remained on the patch for an average duration of 600 ± 120 seconds (*t*_flower-stay_, mean ± SD), simulating a bee visiting a small group of flowers of the same species rather than an individual flower. Agents searched for food sources until they were full, after which they returned to the nest for unloading. They stayed in the nest for 300 seconds (*t*_nest-stay_) (39). The speed of agents moving inside the nest (*v*_nest_) was 0.1 (patch/sec), which allowed them to encounter and recruit other agents (see below).

### Food sources

Either twelve (species-rich environment) or four (species-poor environment) different flower species were in the environment. Foragers needed to visit either 2 (large rewards) or 10 (small rewards) food sources to fill up. In species-rich environments, the number of food sources (*FS*_number_) per flower species was a random number between 0 and 200 (low abundance; *FS*_numberLow_, mean =100 per species) or between 0 and 2000 (high abundance; *FS*_numberHigh_, mean =1000/species). In species-poor environments, it was a random number between 0-600 (low abundance; *FS*_numberLow_, mean =300/species) or between 0 and 6000 (high abundance; *FS*_numberHigh_, mean =3000/species). Thus, flower species-rich and species-poor environments differed in the number of flower species, but not in the number of food sources. The distribution of food sources in the environment was either uniformly random or clustered (Fig. S1a,b). When food sources were clustered, we simulated 10 clusters per flower species (default) at a moderate clustering strength (see Fig. S1b). We also tested environments with 30 clusters, *i.e.* clusters were 3-times more numerous but smaller.

To test if habitat fragmentation affects forage diversity, we simulated a fragmented environment that contained four or eight “build-up” areas, comprising 36% and 50% of the total foraging area, where no food sources were available (thus, 36% or 50% fewer food sources, respectively). This could represent areas with buildings, empty crop fields or roads (Fig. S1c,d).

We tested different refill times (*t*_refill_) for food sources after visits: 0, 1200 (default) and 3600 seconds (75). When *t*_refill_= 0, food sources became rewarding again immediately after the visit of a bee. With *t*_refill_= 1200, a food source remained unrewarding for the equivalent of 20 minutes after it had been visited by a bee, leading to exploitation competition between nestmates.

### Flower constancy

The degree of flower constancy of bees during foraging trips depends on a range of extrinsic and intrinsic factors in nature (7, 9), but is close to 100% in some eusocial bee species (40, 48, 76). In our model, bees visited food sources either indiscriminately (random choice) or they were strictly flower constant, *i.e.* they remained faithful to the flower species they discovered first on their initial foraging trip.

### Recruitment communication

Social bees use different behavioural mechanisms to transmit information about high-quality food sources, such as their odour or location, and thereby bias the food source preferences of their nestmates towards more profitable options (52–55, 77). The model simulated a generic process that allows foragers that have visited a high-quality flower species to bias the food preferences of nestmates during encounters inside the nest (*influencers*). To this end, 3 out of 12 (species-rich environment) or 1 out of 4 (species-poor environment) flower species were designated to be of higher-quality, and foragers visiting these high-quality flower species could become *influencers* upon return to the nest. *Influencers* recruited other agents that were not flower-constant to the high-quality flower species by changing the latter’s preference if they encountered them on the same patch inside the nest. Following such an encounter, recruited agents would leave the nest to search for food sources of the corresponding high-quality flower species. Since the motivation to show communication behaviours often decreases with increasing food source distance (55, 70), the probability of becoming an *influencer* decreased with increasing distance of the last visited high-quality food source (Fig. S3a).

#### Measured variables

To assess how the different parameters affect the dietary diversity of a colony, we measured the total number of flower species visited by the foragers of a colony during a simulation (*Species diversity*). Since forage diversity also depends on the relative abundance (or evenness) of visits, we also calculated the *Simpson’s diversity index* (*SDI*) to assess how evenly the different flower species were collected by a colony. The *SDI* was measured as 1 – *D*, where: 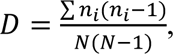 with *n_i_* = the number of food sources in the *i*-th flower species and *N* = the total number of food sources. The *SDI* varies between 0 and 1, with a higher score indicating a higher diversity and evenness of visited flower species.

We tested if *Species diversity* and the *SDI* depended on flower constancy (*vs*. indiscriminate choice), colony size, communication (*vs*. no communication), the total abundance of food sources, and their distribution in flower species-rich and species-poor environments. We performed 10 runs for each parameter combination. We did not calculate statistical *p*-values due to the arbitrariness of the simulation number but indicate 95%- confidence intervals for effect sizes.

### Sensitivity analysis and model exploration

In addition to the factors mentioned above, we explored several other factors and how they affected our results. These included Lévy flight *μ*, refill time, relative reward sizes (larger for high-quality flower species) or the number of high-quality flower species (Table S1).

### Pollen load purity data from the literature

We searched the published literature for information on the strength of flower constancy in different bee species (relying mainly on 18, 35, 44, 47, 75). We included a species if at least 20 bees were sampled and the proportion of bees that collected pure pollen loads (>97% pollen grains belonging to one species) was reported (Table S2). We analysed the effects of lifestyle (as classified by Michener (4): “highly eusocial” = perennial lifestyle with extensive morphological differences between queen and workers; “primitively eusocial” = annual lifestyle and only a moderate morphological difference between queen and workers; and solitary) on flower constancy using phylogenetic generalised least-squares models (PGLS) (78). The phylogenetic framework for the PGLS models relied on phylogenetic trees with branch lengths corresponding to geological time, based on trees for Anthophila (79), Andrenidae (80), Meliponini (81), *Melipona* (82), *Bombus* (83) and *Osmia* (84). A tree was created by pruning species to include only the taxa relevant for the comparative analysis (see Figure S3b).

## Results

### Flower species-rich environment – small rewards

Flower constancy led to a loss of colony forage diversity (Fig. 1a-d), especially in small colonies: when flower constant colonies consisted of 10 foragers, colonies exploited only ∼30-50% of the available plant species (Fig. 1a-d), compared to ∼100% exploited by indiscriminate colonies. As predicted, the percentage of flower species exploited by flower constant bees increased with colony size. When food sources were abundant and clustered (Fig. 1d), however, even large, flower constant colonies (>250 bees) exploited only 60-70% of the available plant species.

**Figure 1.**
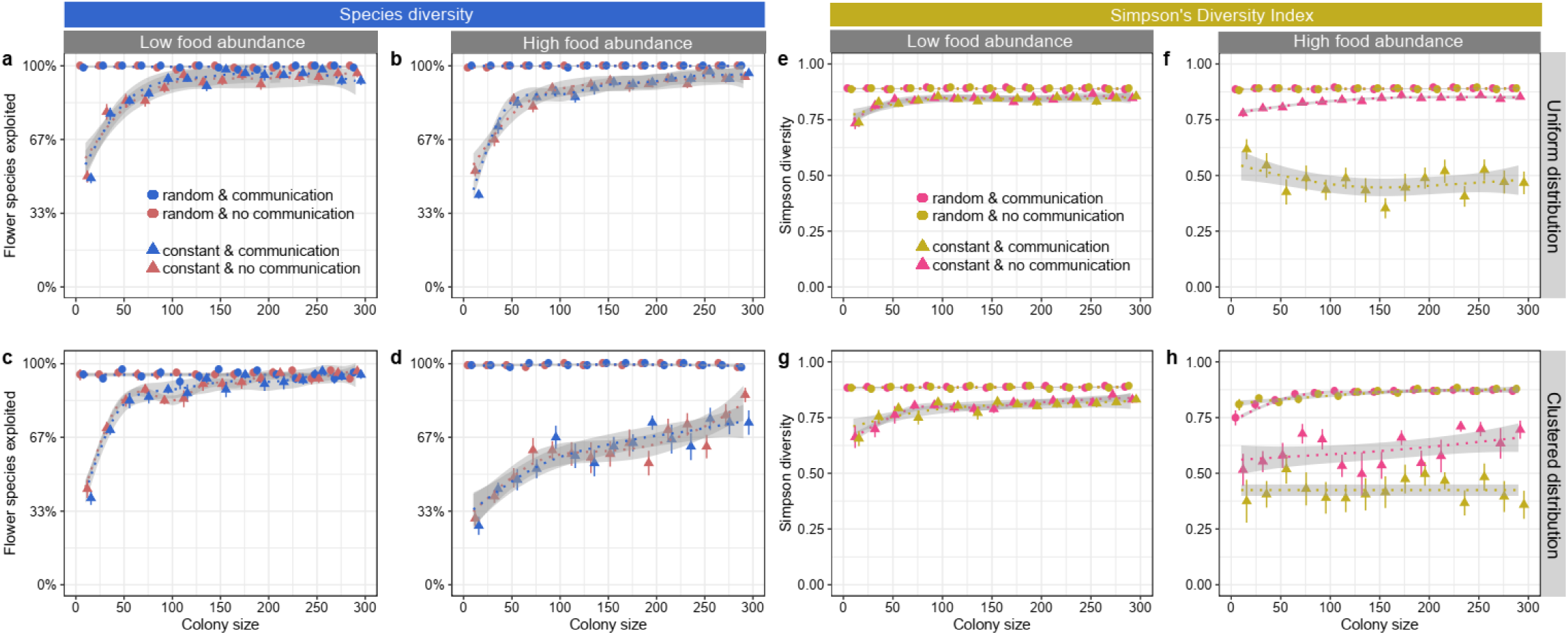
*Species diversity* (as % of all available flower species) (a-d) and *Simpson’s diversity index (SDI)* (e-h) in relation to colony size, food source distribution, food abundance and foraging strategy (flower constancy and communication) when food sources offered *small* rewards and needed time to replenish (1200 seconds). Colonies were either flower constant (triangle) or foraged indiscriminately (= randomly, circles); colonies either had communication (blue in a-d; pink in e-h) or consisted of bees that foraged solitarily (red in a-d; orange in e-h).

The *SDI* was also lower in flower constant colonies compared to indiscriminate colonies (Fig. 1e-h). However, the *SDI* was not greatly affected by colony size. On the other hand, there was a pronounced negative effect of communication on the *SDI* in environments with high food source abundance, even in large colonies (Fig. 1e,g *vs.* 1f,h). For example, when food sources were clustered and colony sizes large (>250 bees), ∼75% of all foraging trips were to just one flower species (Supplementary Information), predominantly a high-quality one.

The reported outcomes refer to food sources that needed time to replenish after a visit (refill = 1200 seconds). We also explored forage diversity when food sources replenished immediately after a visit (refill = 0 seconds), thereby removing exploitation competition. The overall patterns were very similar (Fig. S4).

### Flower species-rich environment – large rewards

When food sources offered large rewards, the negative effects of flower constancy on *Species diversity* were even more pronounced, especially when colonies could also communicate about high-quality flower species (Fig. 2a-d), as increasing colony sizes no longer mitigated against the negative impact of flower constancy. For instance, when food source abundance was high (Fig. 2b,d), flower constant colonies with communication discovered and exploited only ca. 20-30% of the available plant species, irrespective of colony size. The overall patterns were similar when food sources replenished immediately after a visit (Fig. S5).

**Figure 2.**
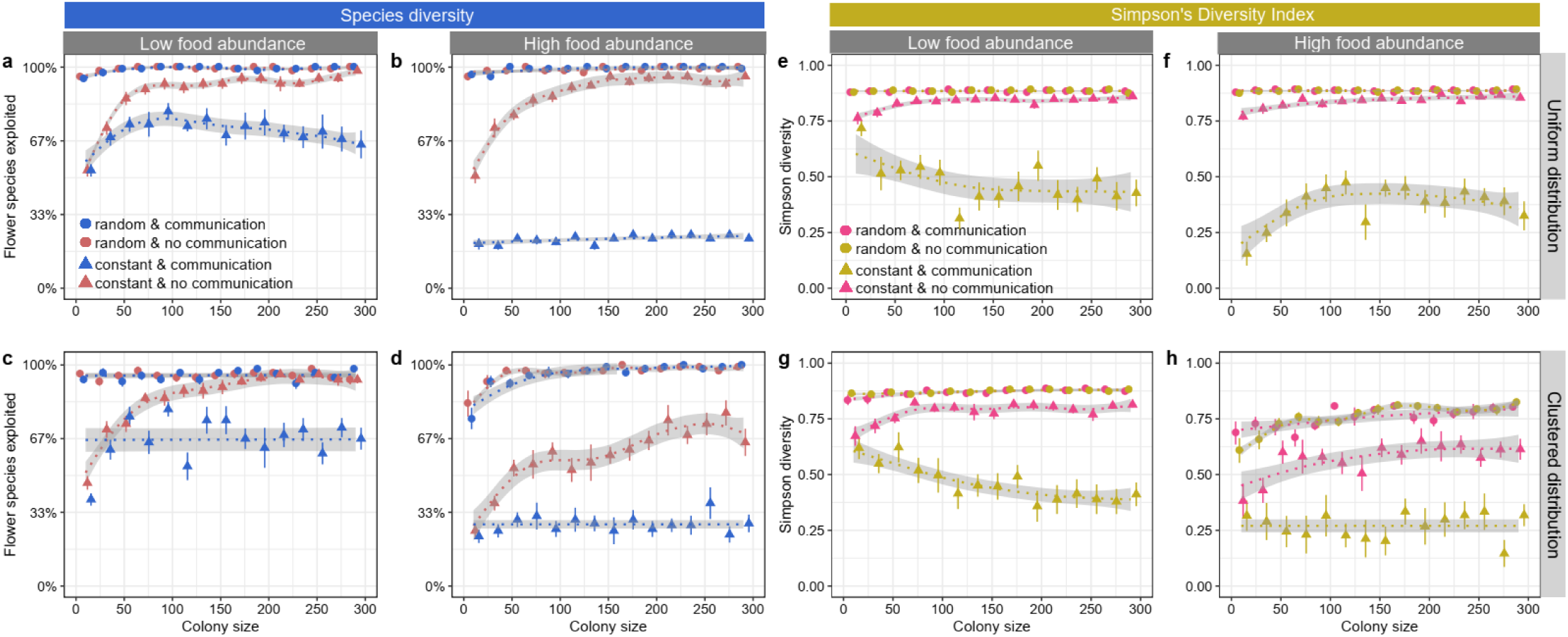
The *Species diversity* (as % of all available flower species) (a-d) and *SDI* (e-h) in relation to colony size, food source distribution, food abundance and foraging strategy (flower constancy and communication) when food sources offered *large* rewards and needed time to replenish (1200 seconds).

The *SDI* was again lower in flower constant colonies (Fig. 2e-h) and communication caused flower constant colonies to focus most of their attention on one or two high-quality plant species. For instance, when food sources were abundant and clustered, ∼84% of all foraging trips of large (>250 bees), flower constant colonies with communication were to just one high-quality plant species (Supplementary Information).

### Flower species-poor environment – small reward

We also explored the effects of flower constancy and communication in environments with low plant diversity (4 plant species). The effects of flower constancy were again similar to what we found in more diverse environments (Fig. 3a-d). However, the effect of colony size was less strong in relative terms: small, flower-constant colonies exploited ca. 70-80% of all available plant species (∼3 flower species), compared to ∼30-50% in a flower species-rich environment (∼4-6 flower species). The loss of forage diversity in flower constant colonies was again more pronounced when food sources were abundant and clustered (Fig. 3c,d).

**Figure 3.**
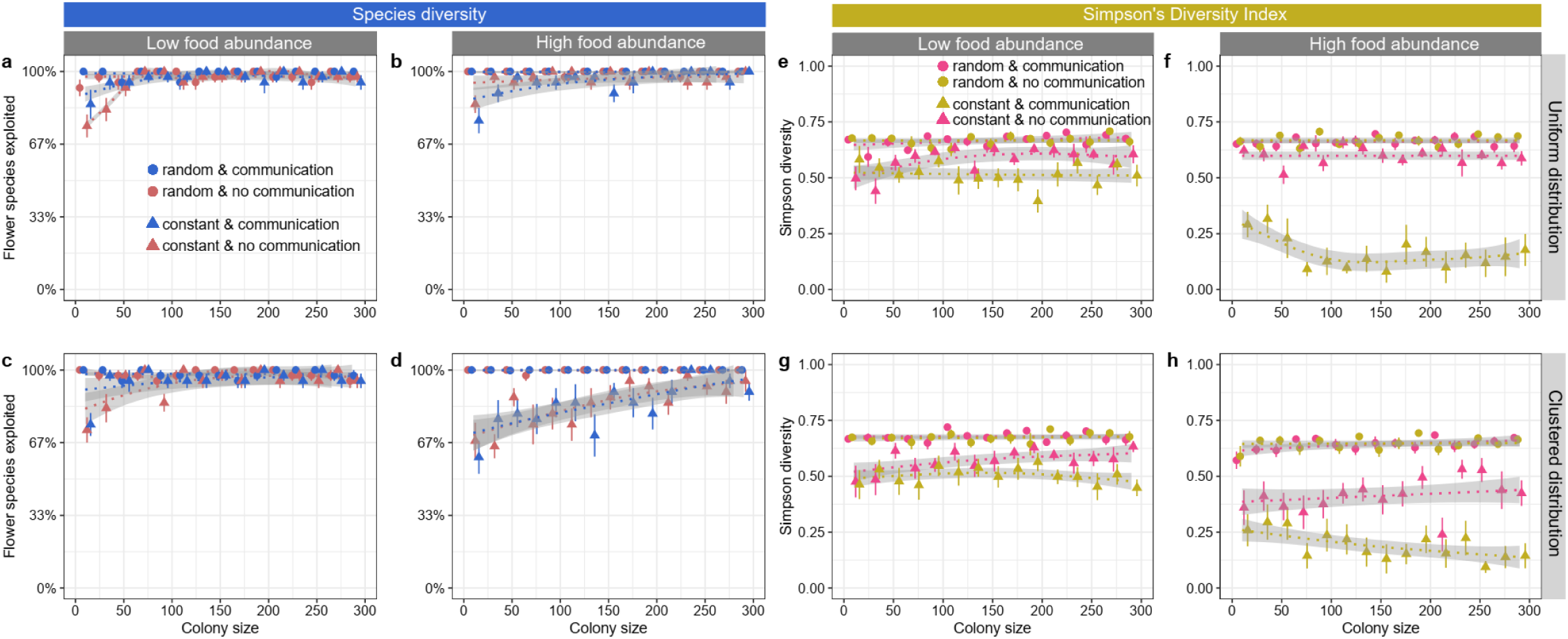
*Species diversity* (as % of all available flower species) (a-d) and *SDI* (e-h) in relation to colony size, food source distribution, food abundance and foraging strategy (flower constancy and communication) in environments with low plant diversity (4 species), when food sources offered *small* rewards and needed time to replenish (1200 seconds).

The *SDI*, on the other hand, showed a contrasting pattern and was generally lower in environments with a low plant diversity (Fig. 3e-h) compared to a flower species-rich environment, revealing a strongly uneven exploitation of food sources. For example, when food sources were clustered, large (>250 foragers) flower constant colonies with communication performed ∼91% of all foraging trips to just one plant species (Supplementary Information). The overall patterns were similar when food sources replenished immediately after a visit (Fig. S6).

### Flower species-poor environment – large rewards

*Species diversity* was again much more impacted by flower constancy and communication when food rewards were large (Fig. 4a-d) compared to when rewards were small (Fig. 3a-d). This effect was especially strong when food source abundance was high (Fig. 4b,d). Under these conditions, flower constant colonies that used communication incorporated only 1-2 of the 4 available flower species into their diet (Fig. 4b,d).

**Figure 4.**
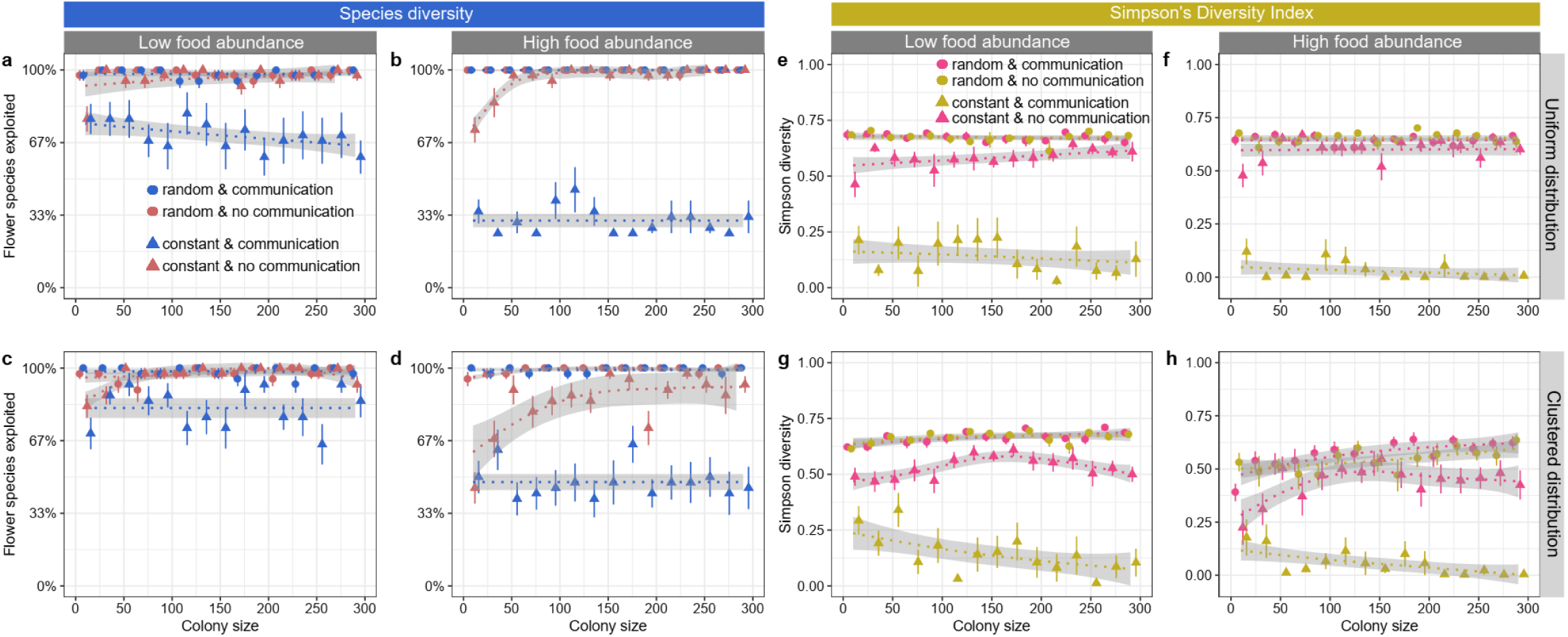
The *Species diversity* (as % of all available flower species) (a-d) and *SDI* (e-h) in relation to colony size, food source distribution, food abundance and foraging strategy (flower constancy and communication). All measurements are from environments with low plant diversity (4 species), in which food sources offered *large* rewards and needed time to replenish (1200 seconds).

The *SDI* approached zero in high food abundance environments and when flower constant colonies also used communication (Fig. 4f, h), meaning that just one flower species was exploited. The overall patterns remained similar when food sources replenished immediately after a visit (Fig. S7).

### Fragmented landscapes and different types of clustering

We simulated colonies in two different fragmented landscapes, mimicking urban and agricultural habitats (Fig. S1c,d). Fragmented environments offered fewer food sources since they were partly covered by squared (covering 36% of the total area) or striped (50% of total area) areas without food. As a result, colonies collected less food in these environments (Supplementary Information; ∼18% and ∼23% fewer food source visits in environments fragmented by squares or stripes, respectively).

We tested if two types of clustering, few large clusters *vs.* many small clusters, differentially affected forage diversity. Unexpectedly, *Species diversity* and the *SDI* were similar in fragmented *vs.* unfragmented environments (Fig. 5). A noteworthy exception is that, when colonies could communicate, large colonies collected food with a lower *SDI* in unfragmented landscapes than in unfragmented landscapes (Fig. 5f, h). Possibly, communication about high-quality flower species was faster in unfragmented environments with more abundant food sources as bees encountered food sources quicker, causing a narrower diet. Forage diversity was overall higher when clusters were smaller but more numerous (30 clusters) than when clusters were larger but fewer in number (10 clusters) (Fig. 5).

**Fig. 5.**
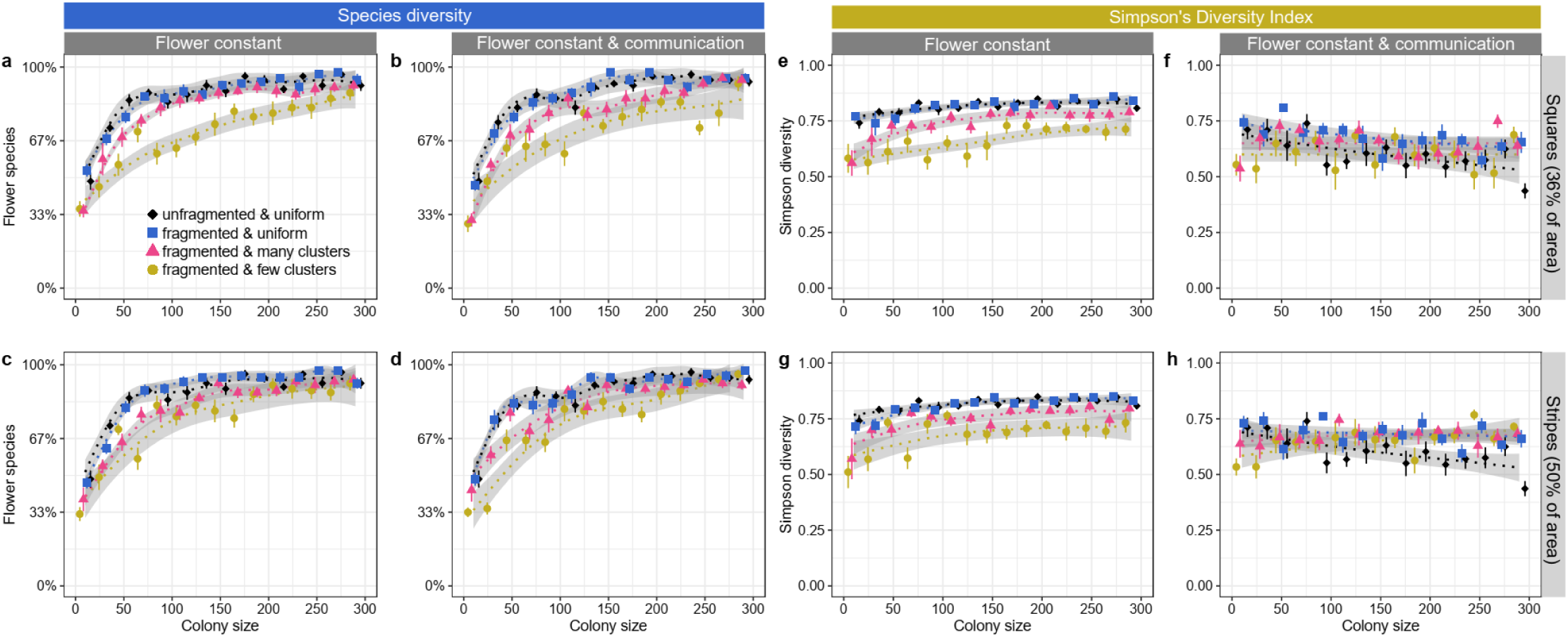
The *Species diversity* (as % of all available flower species) (a-d) and *SDI* (e-h) in relation to colony size, type of fragmentation (squares *vs.* stripes *vs.* unfragmented; see Fig. S1c,d) and foraging strategy (flower constancy and communication). All measurements are from environments with high flower species diversity (12 species), in which uniformly distributed food sources offered *small* rewards and needed time to replenish (1200 seconds). Food source abundance was intermediate in all cases, with an average of 500 food sources per plant species (∼6000 in total).

### Sensitivity analysis and model exploration

In addition to the parameters discussed above, we tested the effects of several other factors and parameters, including food source number (maximum of 50, 100, 500, 1000 per plant species) and longer refill times (3600 seconds). We also tested environments where only one flower species was of high quality and situations when high-quality flower species offered more voluminous rewards than low-quality flower species (50% *vs.* 10% of the load capacity) and we simulated alternative Lévy-flight *μ* values (to 1.4 and 2.4). The general patterns were similar (data can be found in the Supplementary Information). A previous version of the model found that changing recruitment parameters (Fig. S3a) had no noticeable effect on the visitation rates of different flower species (14) and, therefore, these were not explored.

### Pollen load purity

Our comparative analysis included 30 bee species belonging to five bee families (Table S2). Foragers of highly eusocial species (Apini and Meliponini) were more likely to collect pure pollen loads (96.6 ± 3.5% pure loads, N = 12 species) than primitively eusocial species (*Bombus* and a *Lasioglossum*; 57.1 ± 19.3%, N = 8 species) (PGLS: *t* = -2.3, *p* = 0.029) and solitary species (40.6 ± 25.3%, N = 10 species) (*t* = -2.59, *p* = 0.015), suggesting that highly eusocial bee species have the highest degree of flower constancy. There was no significant difference between primitively eusocial and solitary species in how often bees collected pure pollen loads (*t* = -0.8, *p* = 0.43) (Fig. 6; Table S2).

**Fig. 6.**
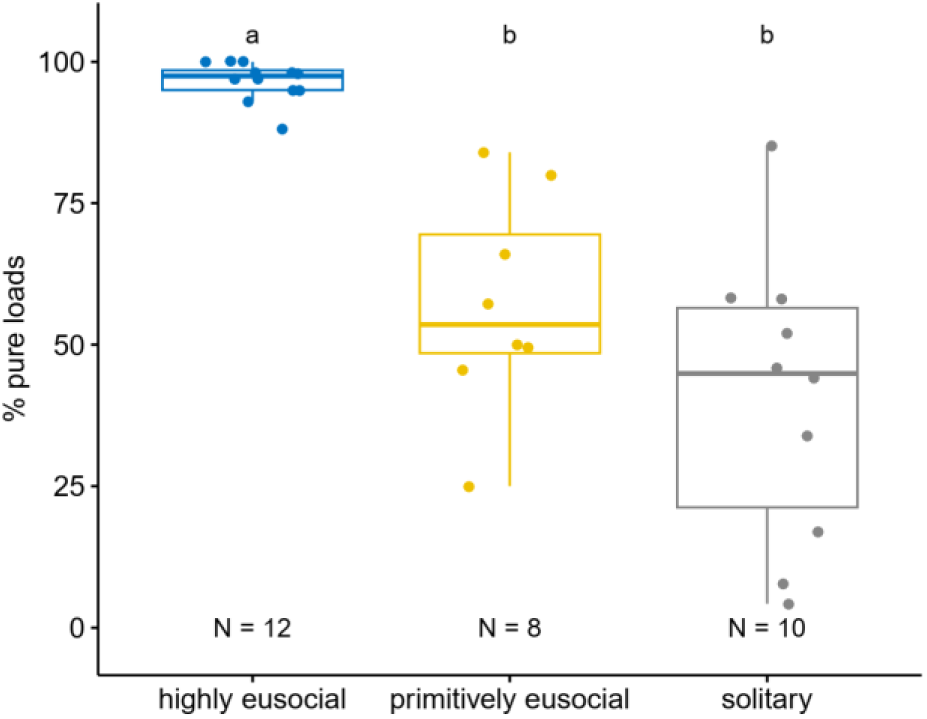
Percentage of foragers collecting pure pollen loads in 30 bee species. Letters a and b indicate statistically significant differences (see Table S2 for details).

## Discussion

Flower constant colonies discovered and exploited only a subset of the available flower species under most conditions, unlike colonies with indiscriminate foragers which exploited close to 100% of all available flower species. Especially small, flower constant colonies often exploited only 30-50% of the available flower species. The *Simpson’s diversity index* (*SDI)*, a measure of how evenly colonies exploited different flower species, was particularly low if flower constant colonies also used communication about high-quality flower species, even when colony sizes were large. Under these conditions, colonies visited predominantly just one or two flower species of higher quality. This is consistent with observations in *Scaptotrigona* stingless bees, which have large colony sizes and an efficient recruitment communication (85, 86): Ramalho (51) observed that while colonies collected pollen from a range of plant species, they mainly concentrated on *Eucalyptus* pollen. Similarly, honey bee (*Apis mellifera*) colonies have been observed to concentrate on highly profitable plant species even if patches are found at distances of several km (87). The ability to focus on profitable flower species is a crucial benefit of communication (52, 55, 59), but it could jeopardise a diverse diet. Collecting only a small number of pollen types increases the risk of missing nutritional intake targets, *e.g.* by collecting pollen with a low protein content (24, 28, 32, 36, 88) or collecting large quantities of pollen containing harmful compounds, such as toxic phytochemicals (35). On the other hand, the high *Species diversity* found in larger flower constant colonies suggests that colony size can help colonies avoid deficiencies in micronutrients, *i.e.* nutrients that are required only in small quantities, even if *Simpson’s diversity* is low. Whether plant *Species diversity* or *Simpson’s diversity* is a more relevant measure of forage diversity is likely to depend on the nutrient type and whether bees require large or small amounts of particular nutrients, which remains unknown for most bee species (23, 31).

Colonies with only 10 foragers discovered only a fraction of the available flower species. Temperate bumble bee colonies have a handful of pollen foragers in spring and even a mature colony of buff-tailed bumble bees (*Bombus terrestris*) or common carder bees (*Bombus pascuorum*) with 100-150 workers (71, 89) contains a relatively small number of pollen foragers (ca. 15-20% of the colony) (39, 91). Many highly eusocial bees, like honey bees or stingless bees, will have several hundred to a few thousand active foragers (54, 70) and are, therefore, predicted to discover a larger number of flower species (*Species diversity*). A higher risk of missing out on key nutrients might explain why bumblebees and solitary bees are less flower constant than honey bees and stingless bees (Fig. 6; 43, 45) and, in turn, why bumble bee colonies often collect pollen from a larger number of flower species than honey bee colonies in the same area (48, 91). Differences in the risk of an unbalanced nutrient intake could also explain why bumble bees, but not honey bees, can discriminate between pollen types based on protein and lipid content (92–95). Honey bees and stingless bees, on the other hand, store pollen inside the nest for longer time periods, which could help bees access a larger diversity of pollen types kept in storage (96) and could allow nurse bees to actively mix pollen types during the preparation of brood food. Whether and how nurse bees that prepare brood food mix stored pollen to create a balanced diet is an intriguing unanswered question.

Simulations suggest that the effects of flower constancy and communication on forage diversity also depend on ecological factors, such as the distribution and abundance of food sources. Paradoxically, forage diversity was often lower when food sources were more abundant (see e.g. Fig. 1). Increased food source abundance means that food sources are discovered quicker, potentially “locking” colonies with flower constancy and communication into collecting certain types of food more rapidly. This effect was amplified by food source clustering, which could mimic large patches or fields of a flower species (see Fig. 1d, 2d and 3d). Food source clusters in nest proximity are likely to be discovered first, which will bias the foraging preferences of a flower constant colony towards this flower species.

When rewards were large, colonies with both flower constancy and communication experienced extremely low forage diversity, both in *Species diversity* and in the *SDI*. One explanation is that foragers collecting larger rewards needed to visit fewer food sources to fill up (e.g. the pollen basket), thereby reducing the amount of time spent searching for food sources and increasing the time spent inside their nest. This is likely to amplify the effects of communication by increasing opportunities for *influencers* to bias the foraging preferences of nestmates towards high-quality flower species, an effect that could be amplified further by the tendency of bees to be more flower constant when the rewards they experience are larger (97).

Humans have drastically modified the environments bees inhabit. Conversion of natural habitat into urban and agricultural spaces has created more fragmented and less species rich habitats (42, 98–100). Reduced floral diversity and the loss of natural habitat are both key drivers of poor nutrition and reduced bee diversity (23, 41–43, 98). Unsurprisingly, forage diversity of flower constant colonies was especially low in species-poor environments in our simulations. The *Simpson’s diversity index* often approached zero as colony size increased and colonies used communication about high-quality flower types (Fig. 4). Flower constancy alone also had a negative impact on the *SDI*, but this effect was less strong in the absence of communication (∼5-20% lower *SDI* compared to indiscriminate colonies) (Fig. 3 and 4). Contrary to our expectation, habitat fragmentation did not reduce forage diversity further in flower constant colonies (Fig. 5; fragmentation reduced the number of food source visits by ∼18-23%, Supplementary Information). We compared two different types of food source clustering, few large clusters *vs.* many smaller clusters, and found that arranging food sources in many smaller clusters increased colony diet breadth by 10-30% compared to environments with fewer but larger clusters. This finding can be tested experimentally and should be considered when planning landscape modifications that aim to improve bee nutrition and health.

In sum, our data suggest that flower constancy is likely to extensively reduce forage diversity in many environments, especially when combined with communication. Flower constant colonies will not only have a lower forage diversity, but they might often collect less food overall as bees skip rewarding food sources that are not of the preferred species (7, 9, 14). This adds further mystery to the question of why social bees are often highly flower constant (Fig. 6) (17). A better understanding of the nutritional needs of bee species is needed to understand the choices and challenges that different bee species face in different environments (23). Furthermore, research is needed to explore the sensory abilities of bees to assess pollen quality and identity, which would allow nurse bees and foragers to follow strategies that reduce the risks of nutritional imbalances (34, 92–95).

## Acknowledgements

We thank Harry Siviter for helpful comments on drafts of this manuscript. This work was carried out using the computational facilities of the Advanced Computing Research Centre, University of Bristol (bristol.ac.uk/acrc).

## Supplementary figures

**Fig. S1.**
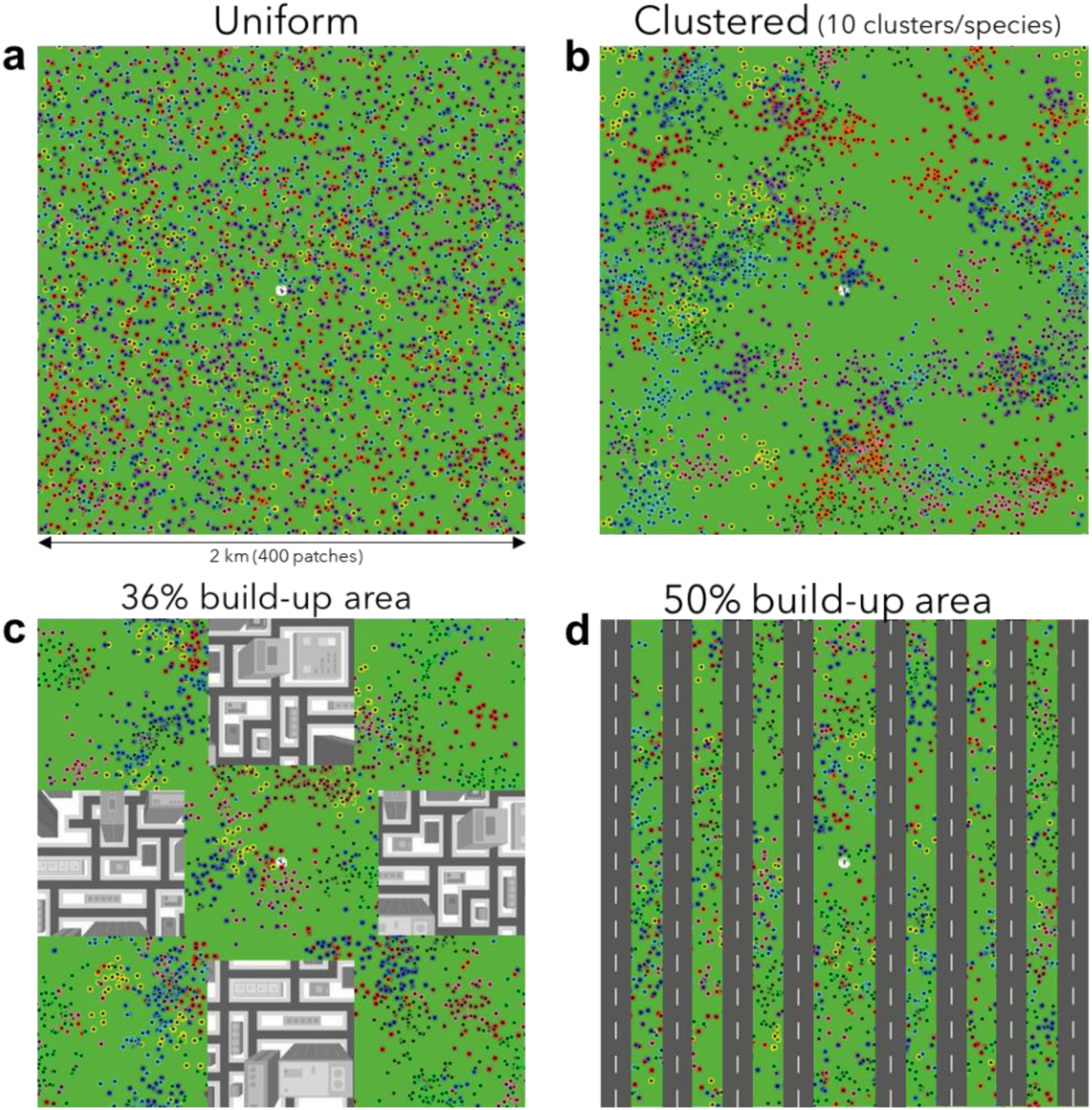
Different types of landscapes simulated in our model. Nests are located in the centre (white circle). Most simulations were done with either uniform (a) or a clustered (b) distribution of food sources. We also explored forage diversity in two different types of fragmented landscapes, with “build-up” area covering either 36% (c) or 50% (d).

**Fig. S2.**
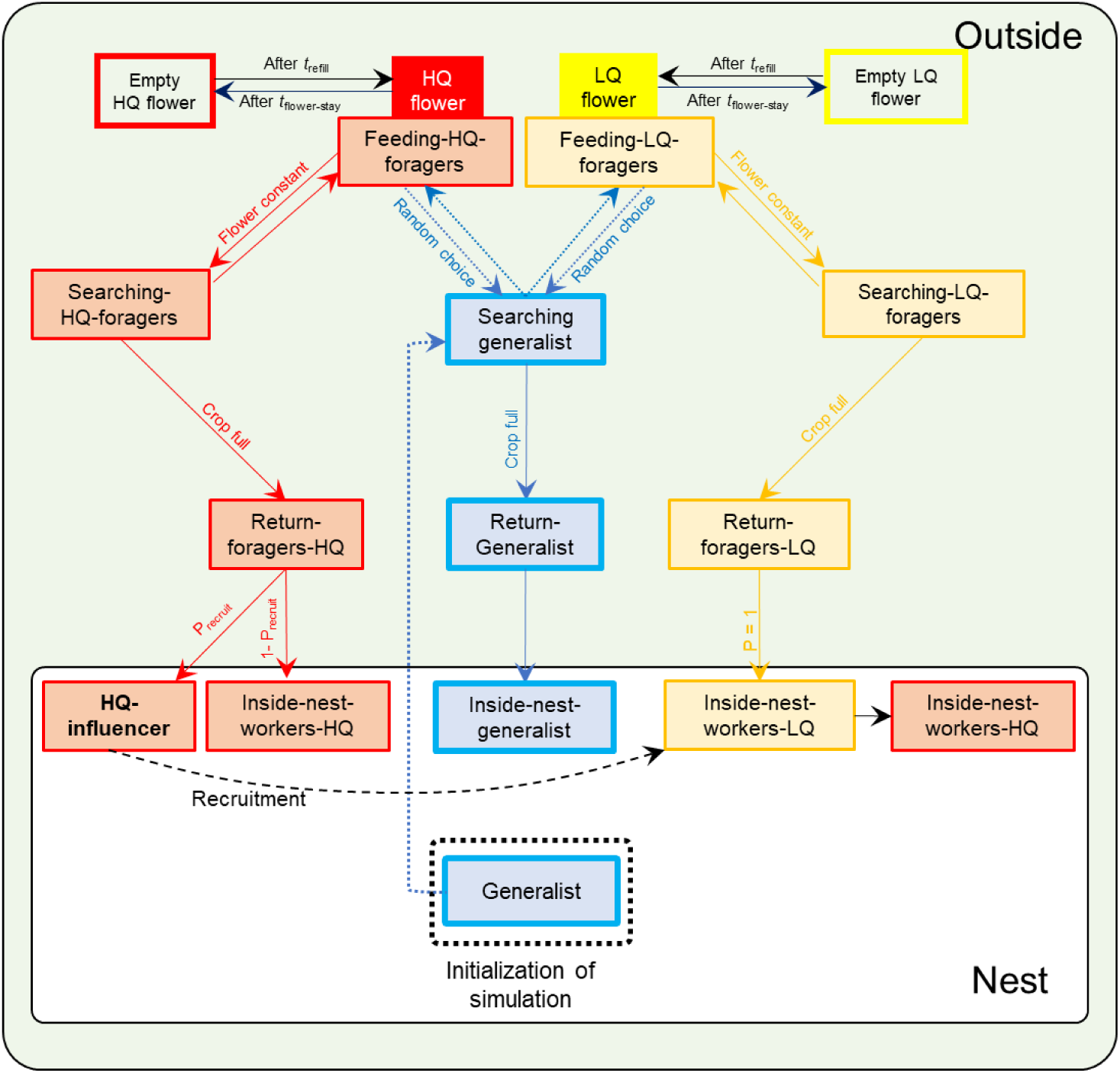
State diagram showing the different states of the agents and the possible transitions between states. Here, LQ flowers represent a lower-quality flower species while HQ flowers represent a high-quality flower species. Flower constant agents visiting high-quality flower species could become *influencers* after their return to the nest and affect the flower preferences of foragers that had previously visited low-quality flower species (see text for details).

**Fig. S3.**
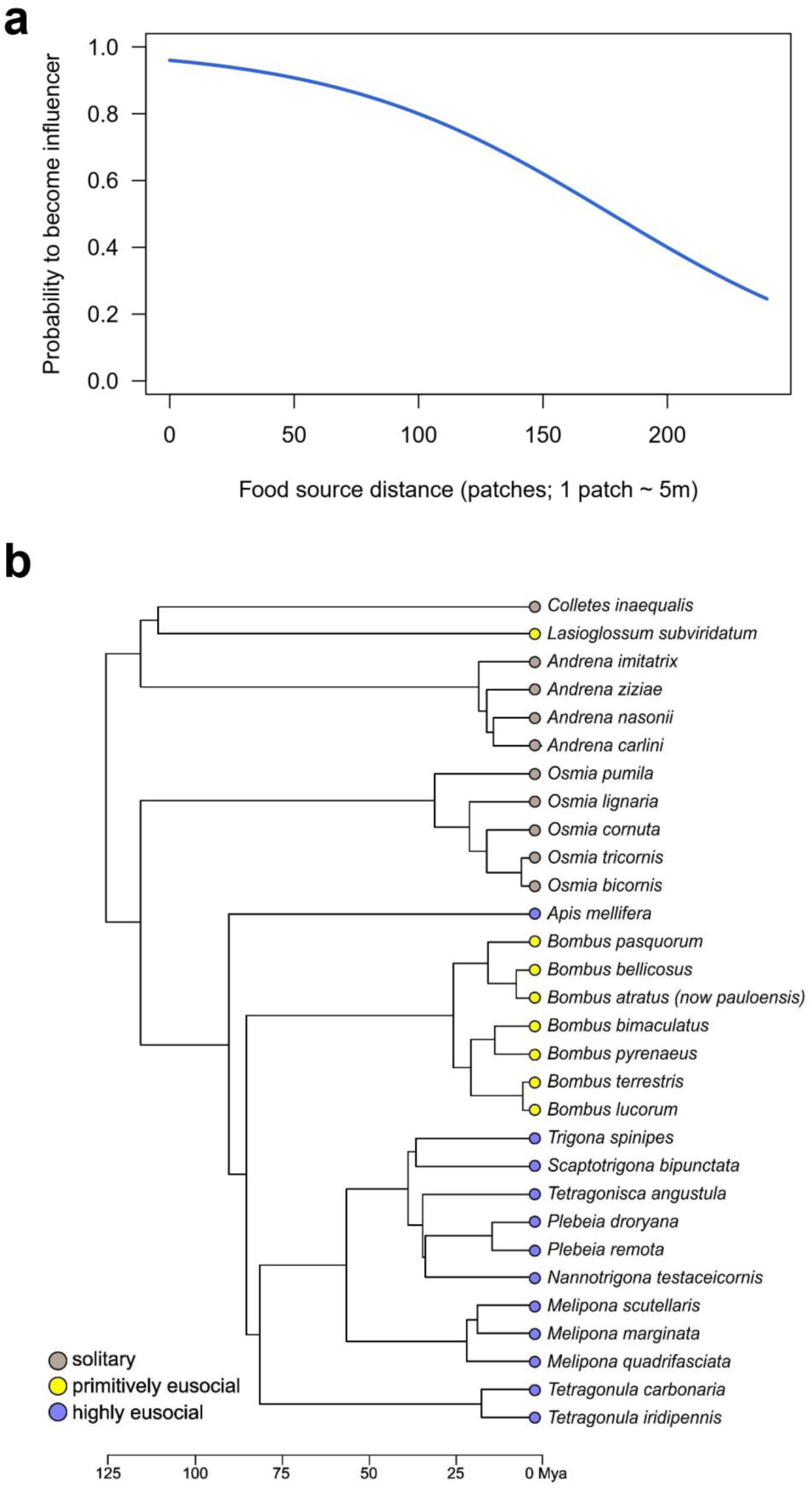
(a) Probability that a returning bee that visited the high-quality flower species becomes an influencer inside the nest. Bees visiting the low-quality flower species did not become influencers under default conditions. Other functions were explored by (14) but they had no noticeable effect on the visitation rates of different flower species (see their Fig. S1 and S3). (b) Pruned phylogenetic tree used for the analysis of the degree of flower constancy (% of foragers collecting pure pollen). Mya = million years ago.

**Figure S4.**
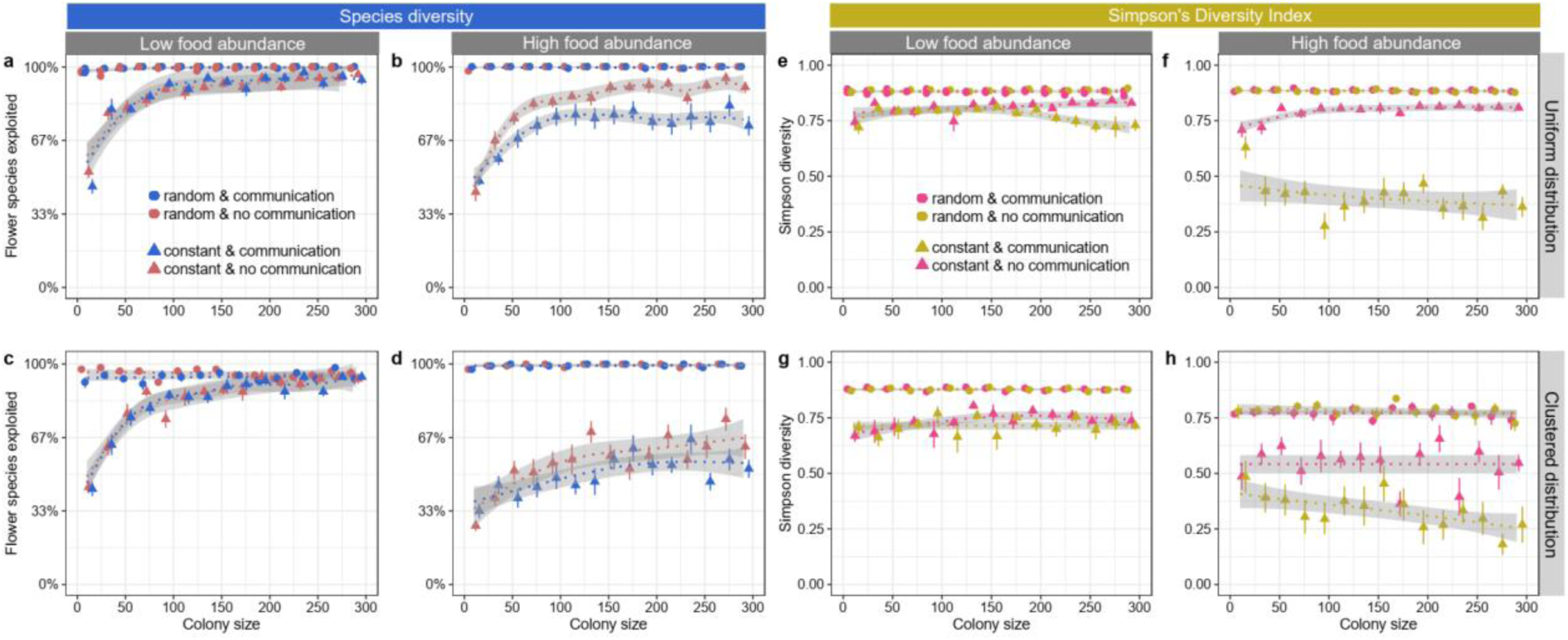
The *Species diversity* (as % of all available flower species) (a-d) and *SDI* (e-h) in relation to colony size, food source distribution, food abundance and foraging strategy (flower constancy and communication). All measurements are from environments in which food sources offered *small* rewards and replenished immediately after a visit. Colonies were either flower constant (triangle) or foraged indiscriminately (circles); bees could use communication (blue in a-d; pink in e-h) or not (red in a-d; orange in e-h).

**Figure S5.**
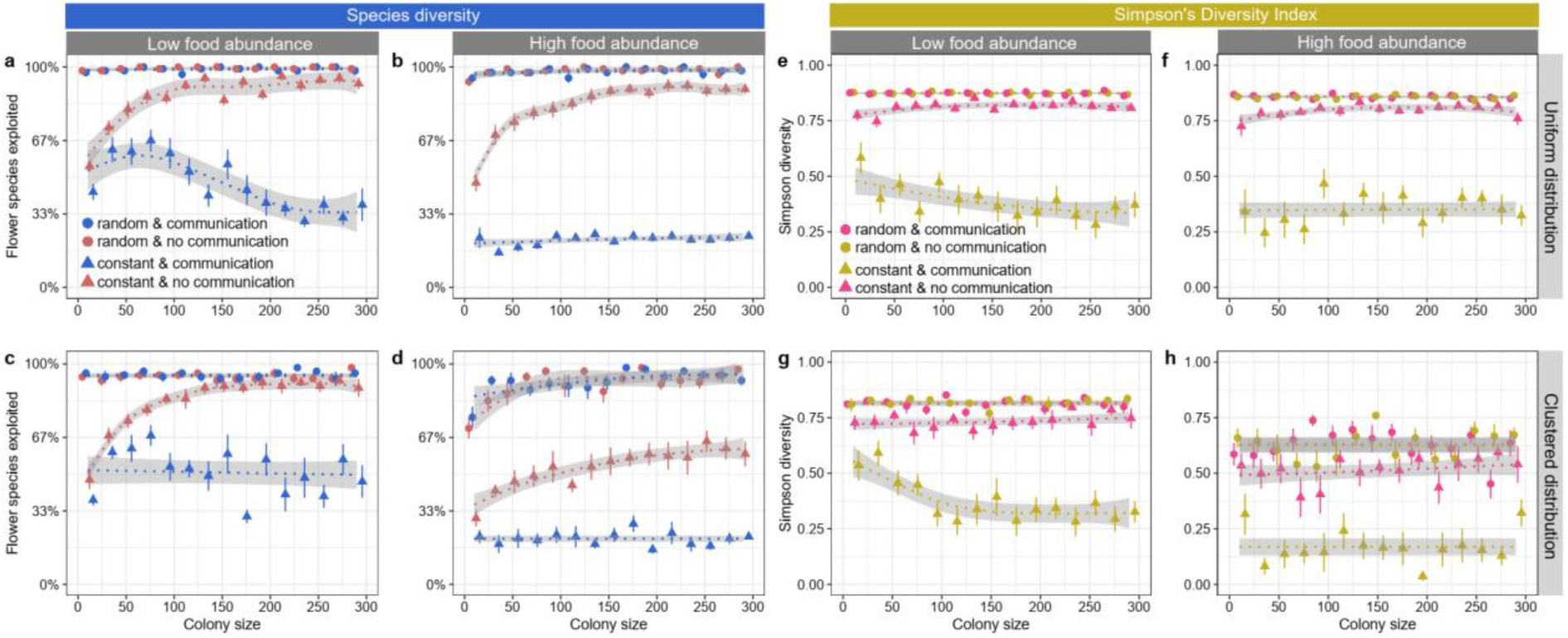
The *Species diversity* (as % of all available flower species) (a-d) and *SDI* (e-h) in relation to colony size, food source distribution, food abundance and foraging strategy (flower constancy and communication) when food sources offered *large* rewards and replenished immediately after a visit.

**Figure S6.**
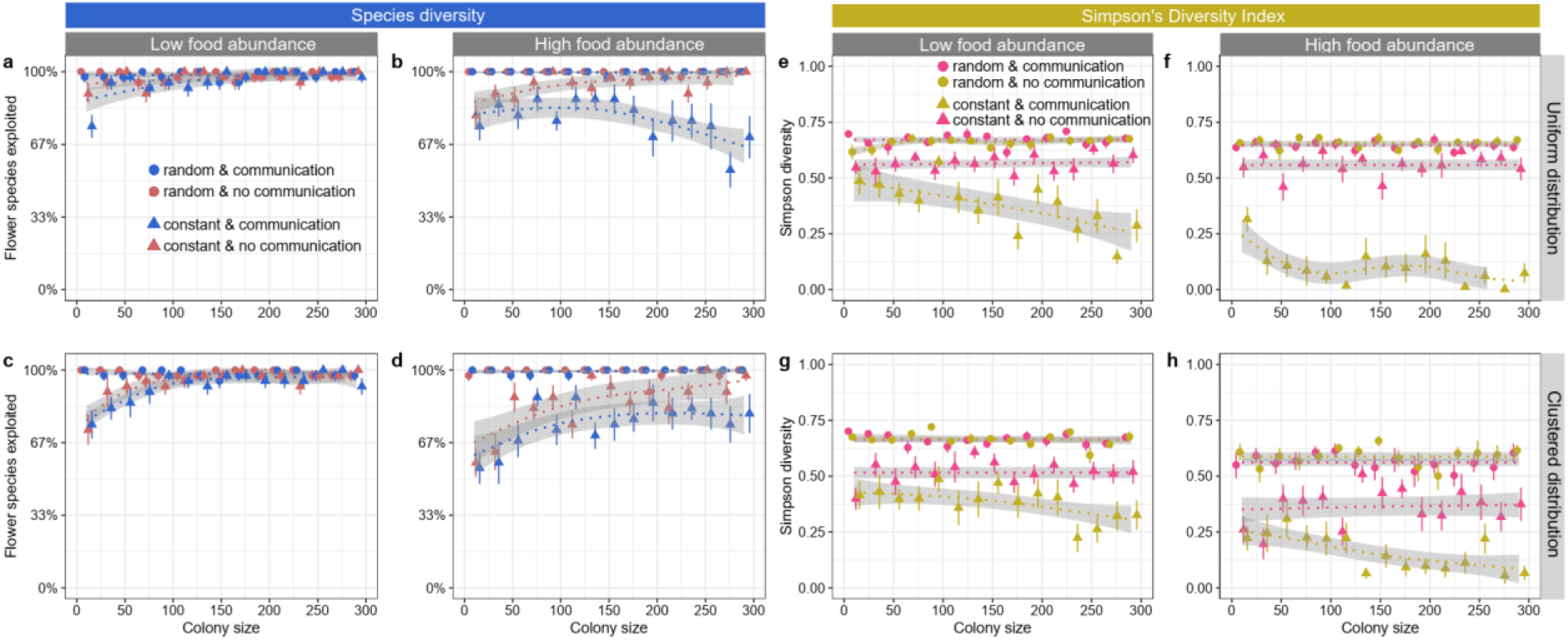
The *Species diversity* (as % of all available flower species) (a-d) and *SDI* (e-h) in relation to colony size, food source distribution, food abundance and foraging strategy (flower constancy and communication) in environments with low plant diversity (4 species), when food sources offer *small* rewards and replenished immediately after a visit (0 seconds).

**Figure S7.**
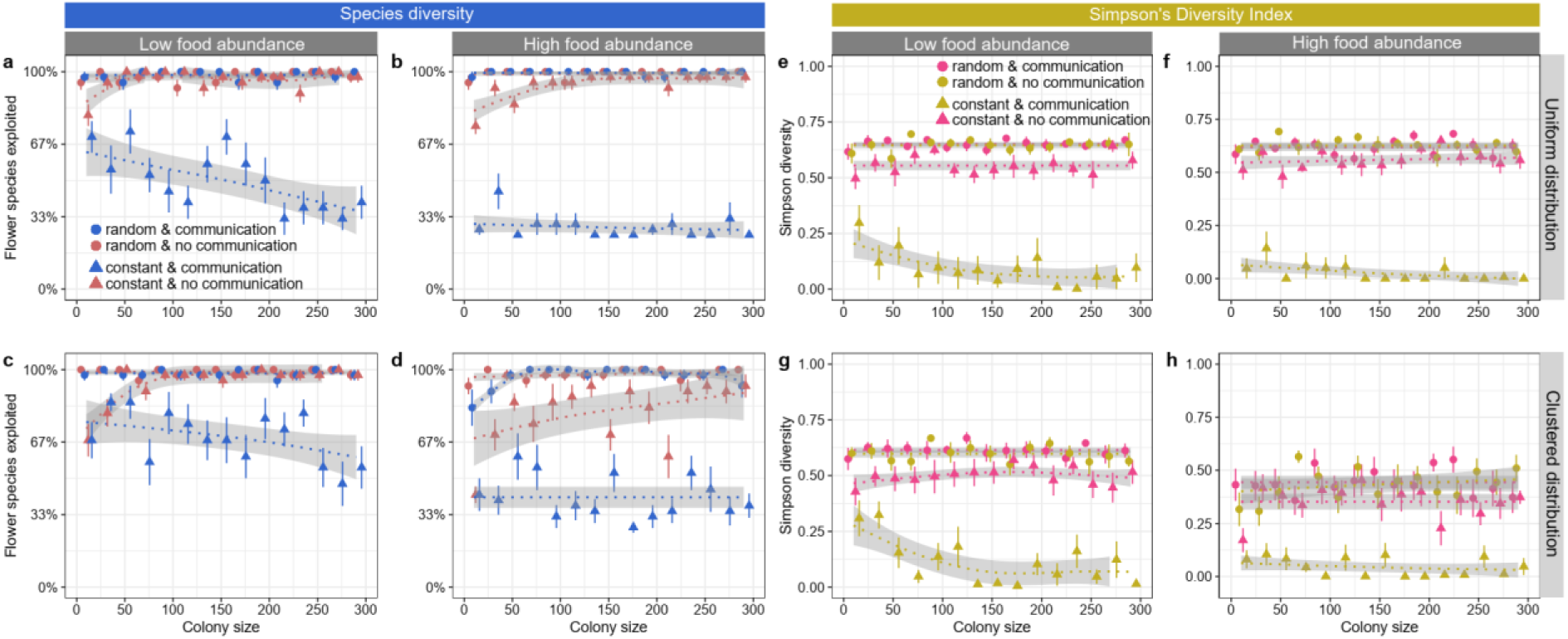
The *Species diversity* (as % of all available flower species) (a-d) and *SDI* (e-h) in relation to colony size, food source distribution, food abundance and foraging strategy (flower constancy and communication) in environments with low plant diversity (4 species), when food sources offer *large* rewards and replenish immediately after a visit (0 seconds).

## Notes

https://doi.org/10.5281/zenodo.8320942

### Competing Interest Statement

The authors have declared no competing interest.

## References

1. K. S. Bawa, Plant-pollinator interactions in Tropical rain forests. Annual Review of Ecology and Systematics 21, 399–422 (1990).

2. A.-M. Klein, et al., Importance of pollinators in changing landscapes for world crops. Proceedings of the Royal Society B: Biological Sciences 274, 303–313 (2007).

3. J. Ollerton, R. Winfree, S. Tarrant, How many flowering plants are pollinated by animals? Oikos 120, 321–326 (2011).

4. C. D. Michener, The bees of the world, 2nd Ed. (The Johns Hopkins University Press, 2007).

5. P. Willmer, Pollination and Floral Ecology (Princeton University Press, 2011).

6. A. J. Bateman, The taxonomic discrimination of bees. Heredity 5, 271–278 (1951).

7. L. Chittka, J. D. Thomson, N. M. Waser, Flower constancy, insect psychology, and plant evolution. Naturwissenschaften 86, 361–377 (1999).

8. C. Darwin, Cross and self fertilization in the vegetable kingdom (Murray, 1876).

9. C. Grüter, F. L. W. Ratnieks, Flower constancy in insect pollinators: Adaptive foraging behaviour or cognitive limitation? Communicative and Integrative Biology 4, 633–636 (2011).

10. N. M. Waser, Flower constancy: definition, cause, and measurement. American Naturalist 127, 593–603 (1986).

11. T.-L. Ashman, G. Arceo-Gómez, Toward a predictive understanding of the fitness costs of heterospecific pollen receipt and its importance in co-flowering communities. American Journal of Botany 100, 1061–1070 (2013).

12. D. R. Campbell, A. F. Motten, The Mechanism of Competition for Pollination between Two Forest Herbs. Ecology 66, 554–563 (1985).

13. C. L. Morales, A. Traveset, Interspecific Pollen Transfer: Magnitude, Prevalence and Consequences for Plant Fitness. Critical Reviews in Plant Sciences 27, 221–238 (2008).

14. L. Hayes, C. Grüter, When should bees be flower constant? An agent-based model highlights the importance of social information and foraging conditions. Journal of Animal Ecology 92, 580–593 (2023).

15. B. Heinrich, “Majoring” and “Minoring” by Foraging Bumblebees, *Bombus vagans*: An Experimental Analysis. Ecology 60, 246–255 (1979).

16. N. E. Raine, L. Chittka, Pollen foraging: learning a complex motor skill by bumblebees (*Bombus terrestris*). Naturwissenschaften 94, 459–464 (2007).

17. C. Smith, L. Weinman, J. Gibbs, R. Winfree, Specialist foragers in forest bee communities are small, social or emerge early. Journal of Animal Ecology 88, 1158–1167 (2019).

18. H. Wells, R. R. S. Rathore, Foraging ecology of the Asian hive bee, *Apis cerana indica*, within artificial flower patches. Journal of Apicultural Research 33, 219–230 (1994).

19. I. Keller, P. Fluri, A. Imdorf, Pollen nutrition and colony development in honey bees: part 1. Bee World 86, 3–10 (2005).

20. R. Brodschneider, K. Crailsheim, Nutrition and health in honey bees. Apidologie 41, 278–294 (2010).

21. A. D. Vaudo, et al., Pollen Protein: Lipid macronutrient ratios may guide broad patterns of bee species floral preferences. Insects 11, 132 (2020).

22. A. D. Vaudo, J. F. Tooker, C. M. Grozinger, H. M. Patch, Bee nutrition and floral resource restoration. Current Opinion in Insect Science 10, 133–141 (2015).

23. M. A. Parreño, et al., Critical links between biodiversity and health in wild bee conservation. Trends in Ecology & Evolution 37, 309–321 (2022).

24. A. Regali, P. Rasmont, Nouvelles méthodes de test pour l’évaluation du régime alimentaire chez des colonies orphelines de *Bombus terrestris* (L) (Hymenoptera, Apidae). Apidologie 26, 273– 281 (1995).

25. T. H. Roulston, J. H. Cane, The effect of pollen protein concentration on body size in the sweat bee *Lasioglossum zephhyrum* (Hymenotpera: Apiformes). Evolutionary Ecology 16, 49–65 (2002).

26. C. Li, B. Xu, Y. Wang, Q. Feng, W. Yang, Effects of dietary crude protein levels on development, antioxidant status, and total midgut protease activity of honey bee (*Apis mellifera ligustica*). Apidologie 43, 576–586 (2012).

27. S. F. Pernal, R. W. Currie, Pollen quality of fresh and 1-year-old single pollen diets for worker honey bees (*Apis mellifera* L.). Apidologie 31, 387–409 (2000).

28. J. O. Schmidt, S. C. Thoenes, M. D. Levin, Survival of honey bees, *Apis mellifera* (Hymenoptera: Apidae), fed various pollen sources. Annals of the Entomological Society of America 80, 176– 183 (1987).

29. T. H. Roulston, J. H. Cane, S. L. Buchmann, What governs protein content of pollen: pollinator preferences, pollen–pistil interactions, or phylogeny? Ecological Monographs 70, 617–643 (2000).

30. A. G. Dolezal, et al., Interacting stressors matter: diet quality and virus infection in honeybee health. Royal Society Open Science 6, 181803 (2019).

31. M. Filipiak, et al., Ecological stoichiometry of the honeybee: Pollen diversity and adequate species composition are needed to mitigate limitations imposed on the growth and development of bees by pollen quality. PLOS ONE 12, e0183236 (2017).

32. A. Génissel, P. Aupinel, C. Bressac, J.-N. Tasei, C. Chevrier, Influence of pollen origin on performance of *Bombus terrestris* micro-colonies. Entomologia Experimentalis et Applicata 104, 329–336 (2002).

33. K. Foley, G. Fazio, A. B. Jensen, W. O. H. Hughes, Nutritional limitation and resistance to opportunistic *Aspergillus* parasites in honey bee larvae. Journal of Invertebrate Pathology 111, 68–73 (2012).

34. F. A. Ruedenauer, et al., Best be(e) on low fat: linking nutrient perception, regulation and fitness. Ecology Letters 23, 545–554 (2020).

35. M. Eckhardt, M. Haider, S. Dorn, A. Müller, Pollen mixing in pollen generalist solitary bees: a possible strategy to complement or mitigate unfavourable pollen properties? Journal of Animal Ecology 83, 588–597 (2014).

36. J.-N. Tasei, P. Aupinel, Nutritive value of 15 single pollens and pollen mixes tested on larvae produced by bumblebee workers (*Bombus terrestris*, Hymenoptera: Apidae). Apidologie 39, 397–409 (2008).

37. C. Alaux, Ducloz François, Crauser Didier, Le Conte Yves, Diet effects on honeybee immunocompetence. Biology Letters 6, 562–565 (2010).

38. G. Di Pasquale, et al., Influence of pollen nutrition on honey bee health: do pollen quality and diversity matter? PLoS ONE 8, e72016 (2013).

39. B. Heinrich, Bumblebee Economics (Harvard University Press, 1979).

40. J. B. Free, The flower constancy of honeybees. Journal of Animal Ecology 32, 119–131 (1963).

41. B. F. Kaluza, et al., Social bees are fitter in more biodiverse environments. Scientific Reports 8, 1–10 (2018).

42. J. Hemberger, M. S. Crossley, C. Gratton, Historical decrease in agricultural landscape diversity is associated with shifts in bumble bee species occurrence. Ecology Letters 24, 1800–1813 (2021).

43. S. H. Woodard, S. Jha, Wild bee nutritional ecology: predicting pollinator population dynamics, movement, and services from floral resources. Current Opinion in Insect Science 21, 83–90 (2017).

44. D. Kleijn, I. Raemakers, A retrospective analysis of pollen host plant use by stable and declining bumble bee species. Ecology 89, 1811–1823 (2008).

45. N. Rossi, E. Santos, S. Salvarrey, N. Arbulo, C. Invernizzi, Determination of flower constancy in *Bombus atratus* Franklin and *Bombus bellicosus* Smith (Hymenoptera: Apidae) through palynological analysis of nectar and corbicular pollen loads. Neotropical Entomology 44, 546– 552 (2015).

46. J. Yourstone, V. Varadarajan, O. Olsson, Bumblebee flower constancy and pollen diversity over time. Behavioral Ecology 34, 602–612 (2023).

47. N. M. Williams, V. J. Tepedino, Consistent mixing of near and distant resources in foraging bouts by the solitary mason bee *Osmia lignaria*. Behavioral Ecology 14, 141–149 (2003).

48. S. D. Leonhardt, N. Blüthgen, The same, but different: pollen foraging in honeybee and bumblebee colonies. Apidologie 43, 449–464 (2012).

49. R. Brodschneider, et al., CSI Pollen: Diversity of honey bee collected pollen studied by citizen scientists. Insects 12, 987 (2021).

50. D. Obregon, G. Nates-Parra, Floral preference of *Melipona eburnea* Friese (Hymenoptera: Apidae) in a Colombian Andean region. Neotropical Entomology 43, 53–60 (2014).

51. M. Ramalho, Foraging by stingless bees of the genus, *Scaptotrigona* (Apidae, Meliponinae). Journal of Apicultural Research 29, 61–67 (1990).

52. D. A. Alves, et al., Diverse communication strategies in bees as a window into adaptations to an unpredictable world. Proceedings of the National Academy of Sciences of the USA 120, e2219031120 (2023).

53. S. Jarau, M. Hrncir, Food Exploitation by Social Insects: Ecological, Behavioral, and Theoretical Approaches, S. Jarau, M. Hrncir, Eds. (CRC Press, Taylor & Francis Group, 2009).

54. C. D. Michener, The Social Behavior of the Bees (Harvard University Press, 1974).

55. K. von Frisch, The dance language and orientation of bees (Harvard University Press, 1967).

56. R. I’Anson Price, C. Grüter, Why, when and where did honey bee dance communication evolve? Frontiers in Ecology and Evolution 3, 1–7 (2015).

57. W. M. Farina, C. Grüter, P. C. Diaz, Social learning of floral odours within the honeybee hive. Proceedings of the Royal Society of London Series B-Biological Sciences 272, 1923–1928 (2005).

58. A. Dornhaus, L. Chittka, Evolutionary origins of bee dances. Nature 401, 38–38 (1999).

59. T. D. Seeley, The wisdom of the hive: The social physiology of honey bee colonies (Harward University Press, 1995).

60. S. P. Hubbell, L. K. Johnson, Comparative foraging behavior of six stingless bee species exploiting a standardized resource. Ecology 59, 1123–1136 (1978).

61. C. Grüter, L. Hayes, Sociality is a key driver of foraging ranges in bees. Current Biology 32, 5390–5397.e3 (2022).

62. A. Dornhaus, F. Klügl, C. Oechslein, F. Puppe, L. Chittka, Benefits of recruitment in honey bees: effects of ecology and colony size in an individual-based model. Behavioral Ecology 17, 336–344 (2006).

63. M. Beekman, J. B. Lew, Foraging in honeybees - when does it pay to dance? Behavioral Ecology 19, 255–262 (2008).

64. M. J. Couvillon, et al., Caffeinated forage tricks honeybees into increasing foraging and recruitment behaviors. Current Biology 25, 2815–2818 (2015).

65. R. I’Anson Price, N. Dulex, N. Vial, C. Vincent, C. Grüter, Honeybees forage more successfully without the “dance language” in challenging environments. Science Advances 5, eaat0450 (2019).

66. M. Begon, J. L. Harper, C. R. Townsend, Ecology: from individuals to ecosystems, Fourth Edition (Blackwell, 2006).

67. U. Wilensky, NetLogo. http://ccl.northwestern.edu/netlogo/ (1999).

68. S. S. Greenleaf, N. M. Williams, R. Winfree, C. Kremen, Bee foraging ranges and their relationship to body size. Oecologia 153, 589–596 (2007).

69. L. K. Kendall, et al., The potential and realized foraging movements of bees are differentially determined by body size and sociality. Ecology 103, e3809 (2023).

70. C. Grüter, Stingless Bees: Their Behaviour, Ecology and Evolution (Springer International Publishing, 2020) (August 23, 2020).

71. C. Westphal, I. Steffan-Dewenter, T. Tscharntke, Bumblebees experience landscapes at different spatial scales: possible implications for coexistence. Oecologia 149, 289–300 (2006).

72. J. D. Crall, S. Ravi, A. M. Mountcastle, S. A. Combes, Bumblebee flight performance in cluttered environments: effects of obstacle orientation, body size and acceleration. Journal of Experimental Biology 218, 2728–2737 (2015).

73. A. M. Reynolds, Lévy flight patterns are predicted to be an emergent property of a bumblebees’ foraging strategy. Behav Ecol Sociobiol 64, 19 (2009).

74. A. M. Reynolds, A. D. Smith, D. R. Reynolds, N. L. Carreck, J. L. Osborne, Honeybees perform optimal scale-free searching flights when attempting to locate a food source. Journal of Experimental Biology 210, 3763–3770 (2007).

75. J. C. Stout, D. Goulson, The Influence of nectar secretion rates on the responses of bumblebees (*Bombus* spp.) to previously visited flowers. Behavioral Ecology and Sociobiology 52, 239–246 (2002).

76. M. Ramalho, T. C. Giannini, K. S. Malagodi-Braga, V. L. Imperatriz-Fonseca, Pollen harvest by stingless bee foragers (Hymenoptera, Apidae, Meliponinae). Grana 33, 239–244 (1994).

77. W. M. Farina, C. Grüter, A. Arenas, “Olfactory information transfer during recruitment in honey bees” in Honeybee Neurobiology and Behavior - A Tribute to Randolf Menzel, C. G. Galizia, D. Eisenhardt, M. Giurfa, Eds. (Springer, 2012), pp. 89–101.

78. E. Paradis, Analysis of Phylogenetics and Evolution with R (Springer Science & Business Media, 2011).

79. S. Cardinal, S. L. Buchmann, A. L. Russell, The evolution of floral sonication, a pollen foraging behavior used by bees (Anthophila). Evolution 72, 590–600 (2018).

80. G. Pisanty, R. Richter, T. Martin, J. Dettman, S. Cardinal, Molecular phylogeny, historical biogeography and revised classification of andrenine bees (Hymenoptera: Andrenidae). Molecular Phylogenetics and Evolution 170, 107151 (2022).

81. C. Rasmussen, S. Cameron, Global stingless bee phylogeny supports ancient divergence, vicariance, and long distance dispersal. Biological Journal of the Linnean Society 99, 206–232 (2010).

82. S. R. Ramírez, et al., A molecular phylogeny of the stingless bee genus *Melipona* (Hymenoptera: Apidae). Molecular Phylogenetics and Evolution 56, 519–525 (2010).

83. M. P. Arbetman, G. Gleiser, C. L. Morales, P. Williams, M. A. Aizen, Global decline of bumblebees is phylogenetically structured and inversely related to species range size and pathogen incidence. Proceedings of the Royal Society B: Biological Sciences 284, 20170204 (2017).

84. M. Haider, S. Dorn, C. Sedivy, A. Müller, Phylogeny and floral hosts of a predominantly pollen generalist group of mason bees (Megachilidae: Osmiini). Biological Journal of the Linnean Society 111, 78–91 (2014).

85. M. Lindauer, W. E. Kerr, Communication between the workers of stingless bees. Bee World 41, 29–71 (1960).

86. F. G. Barth, M. Hrncir, S. Jarau, Signals and cues in the recruitment behavior of stingless bees (Meliponini). Journal of Comparative Physiology A 194, 313–327 (2008).

87. M. Beekman, F. L. W. Ratnieks, Long-range foraging by the honey-bee, *Apis mellifera* L. Functional Ecology 14, 490–496 (2000).

88. M. Vanderplanck, et al., How Does Pollen Chemistry Impact Development and Feeding Behaviour of Polylectic Bees? PLOS ONE 9, e86209 (2014).

89. R. Cueva del Castillo, S. Sanabria-Urbán, M. A. Serrano-Meneses, Trade-offs in the evolution of bumblebee colony and body size: a comparative analysis. Ecology and Evolution 5, 3914–3926 (2015).

90. T. Allen, S. Cameron, R. McGinley, B. Heinrich, The role of workers and new queens in the ergonomics of a bumblebee colony (Hymenoptera: Apoidea). Journal of the Kansas Entomological Society 51, 329–342 (1978).

91. K. K. Graham, et al., Identity and diversity of pollens collected by two managed bee species while in blueberry fields for pollination. Environmental Entomology, in press (2023).

92. A. D. Vaudo, H. M. Patch, D. A. Mortensen, J. F. Tooker, C. M. Grozinger, Macronutrient ratios in pollen shape bumble bee (*Bombus impatiens*) foraging strategies and floral preferences. Proceedings of the National Academy of Sciences of the USA 113, E4035–E4042 (2016).

93. T. K. Kitaoka, J. C. Nieh, Bumble bee pollen foraging regulation: role of pollen quality, storage levels, and odor. Behavioral Ecology and Sociobiology 63, 625–625 (2009).

94. S. F. Pernal, R. W. Currie, The influence of pollen quality on foraging behavior in honeybees (*Apis mellifera* L.). Behavioral Ecology and Sociobiology 51, 53–68 (2001).

95. M. Beekman, K. Preece, T. M. Schaerf, Dancing for their supper: Do honeybees adjust their recruitment dance in response to the protein content of pollen? Insectes Sociaux 63, 117–126 (2016).

96. H. P. Hendriksma, S. Shafir, Honey bee foragers balance colony nutritional deficiencies. Behavioral Ecology and Sociobiology 70, 509–517 (2016).

97. C. Grüter, H. Moore, N. Firmin, H. Helanterä, F. L. W. Ratnieks, Flower constancy in honey bee foragers (*Apis mellifera*) depends on ecologically realistic rewards. Journal of Experimental Biology 214, 1397–1402 (2011).

98. A. G. Dolezal, A. L. St. Clair, G. Zhang, A. L. Toth, M. E. O’Neal, Native habitat mitigates feast– famine conditions faced by honey bees in an agricultural landscape. Proceedings of the National Academy of Sciences of the USA 116, 25147–25155 (2019).

99. C. M. Kennedy, et al., A global quantitative synthesis of local and landscape effects on wild bee pollinators in agroecosystems. Ecology Letters 16, 584–599 (2013).

100. S. G. Potts, et al., Global pollinator declines: trends, impacts and drivers. Trends in Ecology & Evolution 25, 345–353 (2010).

