## Supplemental Table 1 for "Extensive loss of forage diversity in social bees due to flower constancy and communication in simulated environments"

**Table S1:** Overview of the model variables and the used values.

| **Variables** | **Description** | **Default values** | **Other values tested** | **Information source** |
| --- | --- | --- | --- | --- |
| Colony size | Number of foragers in a colony | 10-300 |  | (1, 2) |
| Nest size | The radius of the nest in patches | 5 |  | arbitrary |
| *FS*_numberLow_ | Number of food sources per flower species under low abundance conditions in species-poor and species-rich environments | 0-200, 0-600 | 0-50 and 0-100 | arbitrary |
| *FS*_numberHigh_ | Number of food sources per flower species under high abundance conditions in species-poor and species-rich environments | 0-2000, 0-6000 | 0-500 and 0-1000 | arbitrary |
| *FS* species | Number of flower species | 4 and 12 |  | arbitrary |
| *HQ FS species* | Number of high-quality flower species | 3 | 1 | arbitrary |
| Reward size | Number of food sources that bees needed to visit to fill up | 2, 10 |  | arbitrary |
| *v*_flight_ | Flight speed | 1 patch/tick |  | (3) |
| *t*_flower-stay_ | Time spent at food source, mean ± SD | 600 ± 120 ticks |  | arbitrary |
| *v*_nest_ | Movement speed inside nest | 0.1 patch/tick |  | arbitrary |
| *t*_nest-stay_ | Time in nest between trips | 300 ticks |  | (4) |
| *Clusters* | Number of clusters per flower species when food sources are clustered | 0, 10 | 30 | arbitrary |
| *t*_refill_ | Time until food sources offer food again | 0, 1200 ticks | 3600 | (5) |
| Lévy *μ* | Lévy flight parameter | 1.8 | 1.4, 2.4 | (6) |

**References**

1.	C. Grüter, Stingless Bees: Their Behaviour, Ecology and Evolution (Springer International Publishing, 2020) (August 23, 2020).

2.	C. Westphal, I. Steffan-Dewenter, T. Tscharntke, Bumblebees experience landscapes at different spatial scales: possible implications for coexistence. Oecologia **149**, 289–300 (2006).

3.	J. D. Crall, S. Ravi, A. M. Mountcastle, S. A. Combes, Bumblebee flight performance in cluttered environments: effects of obstacle orientation, body size and acceleration. J. Exp. Biol. **218**, 2728–2737 (2015).

4.	B. Heinrich, Bumblebee Economics (Harvard University Press, 1979).

5.	J. C. Stout, D. Goulson, The Influence of Nectar Secretion Rates on the Responses of Bumblebees (Bombus spp.) to Previously Visited Flowers. Behav. Ecol. Sociobiol. **52**, 239–246 (2002).

6.	A. M. Reynolds, J. L. Swain, A. D. Smith, A. P. Martin, J. L. Osborne, Honeybees use a Lévy flight search strategy and odour-mediated anemotaxis to relocate food sources. Behav. Ecol. Sociobiol. **64**, 115–123 (2009).
