## Supplemental Table 2 for "Extensive loss of forage diversity in social bees due to flower constancy and communication in simulated environments"

**Table S2:** Flower constancy (% of bees collecting pure pollen loads) in 30 bee species.

| **Species** | **Family** | **Type** | **% of bees with pure loads** | **Reference** |
| --- | --- | --- | --- | --- |
| *Tetragonula iridipennis* | Apidae | highly eusocial | 100 | (1) |
| *Tetragonula carbonaria* | Apidae | highly eusocial | 88 | (2) |
| *Melipona quadrifasciata* | Apidae | highly eusocial | 100 | (3) |
| *Melipona marginata* | Apidae | highly eusocial | 98 | (3) |
| *Melipona scutellaris* | Apidae | highly eusocial | 98 | (3) |
| *Nannotrigona testaceicornis* | Apidae | highly eusocial | 100 | (3) |
| *Plebeia remota* | Apidae | highly eusocial | 98 | (3) |
| *Plebeia droryana* | Apidae | highly eusocial | 95 | (3) |
| *Tetragonisca angustula* | Apidae | highly eusocial | 97 | (3) |
| *Scaptotrigona bipunctata* | Apidae | highly eusocial | 93 | (3) |
| *Trigona spinipes* | Apidae | highly eusocial | 95 | (3) |
| *Bombus lucorum* | Apidae | primitively eusocial | 66 | (4) |
| *Bombus terrestris* | Apidae | primitively eusocial | 49.5 | (5, 6) |
| *Bombus pyrenaeus* | Apidae | primitively eusocial | 50 | (7) |
| *Bombus bimaculatus* | Apidae | primitively eusocial | 25 | (8) |
| *Bombus atratus (now pauloensis)* | Apidae | primitively eusocial | 80 | (9) |
| *Bombus bellicosus* | Apidae | primitively eusocial | 84 | (9) |
| *Bombus pasquorum* | Apidae | primitively eusocial | 45.5 | (4, 5) |
| *Apis mellifera* | Apidae | highly eusocial | 97 | (5, 10) |
| *Osmia bicornis* | Megachilidae | solitary | 44 | (11) |
| *Osmia tricornis* | Megachilidae | solitary | 34 | (11) |
| *Osmia cornuta* | Megachilidae | solitary | 52 | (11) |
| *Osmia lignaria* | Megachilidae | solitary | 58 | (11) |
| *Osmia pumila* | Megachilidae | solitary | 17 | (8) |
| *Andrena carlini* | Andrenidae | solitary | 7.7 | (8) |
| *Andrena nasonii* | Andrenidae | solitary | 4.2 | (8) |
| *Andrena ziziae* | Andrenidae | solitary | 85.2 | (8) |
| *Andrena imitatrix* | Andrenidae | solitary | 45.8 | (8) |
| *Lasioglossum subviridatum* | Halictidae | primitively eusocial | 57.1 | (8) |
| *Colletes inaequalis* | Colletidae | solitary | 58.3 | (8) |

**References**

1.	U. Layek, P. Karmakar, Nesting characteristics, floral resources, and foraging activity of Trigona iridipennis Smith in Bankura district of West Bengal, India. *Insect. Soc.* **65**, 117–132 (2018).

2.	D. White, B. W. Cribb, T. A. Heard, Flower constancy of the stingless bee Trigona carbonaria Smith (Hymenoptera: Apidae: Meliponini). *Australian Journal of Entomology* **40**, 61–64 (2001).

3.	M. Ramalho, T. C. Giannini, K. S. Malagodi-Braga, V. L. Imperatriz-Fonseca, Pollen Harvest by Stingless Bee Foragers (Hymenoptera, Apidae, Meliponinae). *Grana* **33**, 239–244 (1994).

4.	J. B. Free, The flower constancy of bumblebees. *Journal of Animal Ecology* **39**, 395–402 (1970).

5.	S. D. Leonhardt, N. Blüthgen, The same, but different: pollen foraging in honeybee and bumblebee colonies. *Apidologie* **43**, 449–464 (2012).

6.	J. Yourstone, V. Varadarajan, O. Olsson, Bumblebee flower constancy and pollen diversity over time. *Behavioral Ecology*, arad028 (2023).

7.	E. Kozuharova, Flower Constancy of Bumblebees – The Case of (Fabaceae) Pollinators. *Journal of Apicultural Science* **62**, 135–140 (2018).

8.	C. Smith, L. Weinman, J. Gibbs, R. Winfree, Specialist foragers in forest bee communities are small, social or emerge early. *Journal of Animal Ecology* **88**, 1158–1167 (2019).

9.	N. Rossi, E. Santos, S. Salvarrey, N. Arbulo, C. Invernizzi, Determination of Flower Constancy in Bombus atratus Franklin and Bombus bellicosus Smith (Hymenoptera: Apidae) through Palynological Analysis of Nectar and Corbicular Pollen Loads. *Neotrop Entomol* **44**, 546–552 (2015).

10.	J. B. Free, The flower constancy of honeybees. *Journal of Animal Ecology* **32**, 119–131 (1963).

11.	M. Eckhardt, M. Haider, S. Dorn, A. Müller, Pollen mixing in pollen generalist solitary bees: a possible strategy to complement or mitigate unfavourable pollen properties? *Journal of Animal Ecology* **83**, 588–597 (2014).
